# Scientific note on the potential beneficial effect of 1-octen-3-ol on the survival of *Apis mellifera* chronically exposed to low doses of fipronil

**DOI:** 10.1101/2024.06.21.599996

**Authors:** Vincent Fernandes, Thania Sbaghdi, Marine Suchet, Frédéric Delbac, Nicolas Blot, Philippe Bouchard

## Abstract

Fipronil, a non-competitive antagonist of γ-aminobutyric acid receptors (GABA-Rs), inhibits the flow of chloride ions in the nerve cells, leading to the insect death. In a previous work, the volatolome of bees chronically exposed to this insecticide (0.5 and 1 *μ*g/L) was analyzed and characterize the *Apis mellifera* metabolic responses. Two Volatile Organic Compounds or VOCs (1-octen-3-ol and 2,6-dimethylcyclohexanol) produced by bee were highlighted. We assumed that these VOCs could act as modulators of the GABA-R to counteract the effect of fipronil. The toxicity of these VOCs was tested on emerging bees for concentrations ranging from 0 to 3.4 *μ*g/L. Survival and the sucrose consumption were recorded during 21 days of chronic exposure. 1-octen-3-ol and 2,6-dimethylcyclohexanol did not significantly affect the proportion of survival at the end of the experiment, whatever their concentration. 1-octen-3-ol (0.5, 1, 1.7 and 3.4 *μ*g/L) was chosen to assess its effect in the case of co-exposure with fipronil (0.5 and 1 *μ*g/L). A beneficial effect on survival at 21 days was observed with an average improvement in survival rate for the co-exposed groups. This positive effect was not related to the VOC concentration and could be a direct effect (interaction with GABA-R) or a hormetic effect (global improvement of bee fitness).

---

The honeybees *Apis mellifera* is a non-target organism of a variety of pesticides (Johnson 2015), including the neurotoxic fipronil a broad-spectrum systemic insecticide (Simon-Delso et al. 2015). Among the responses observed in honeybees exposed to low doses of fipronil, two Volatile Organic Compounds (VOCs) have shown a modulated abundance in the abdomen, the 2,6-dimethylcyclohexanol, hereafter designed as DMCH, and the 1-octen-3-ol, hereafter named OO (Fernandes et al. 2023 and unpublished data). These VOCs have been described as modulators of γ-aminobutyric acid receptors (GABA-R) in mammals (Chowdhury et al. 2016; Johnston 2006; Köksal 2015). The binding of DMCH or OO to the mammalian GABA-R leads to the prolonged opening of the chloride channels. In insects, the GABA-R is the main target of fipronil (Casida and Durkin 2013). Unlike to the VOCs, fipronil binding to GABA-R blocks the chloride ions flow, resulting in neuron hyperexcitability. It is not known whether the VOCs bind to the insect GABA-R and exert opposite effect to that of fipronil, eventually attenuating fipronil-induced hyperexcitation. To test whether VOCs could counterbalance insecticide-induced mortality, honeybees were chronically exposed to fipronil and/or VOCs.

Since VOCs may be toxic to organisms (Leroy et al. 2011; Saha and Chandran 2017), the toxicity of each VOC was assessed on the honeybee survival. Workers were collected on frames from an experimental apiary (Université Clermont Auvergne) in spring 2023. They were mixed and distributed into cages and fed with sucrose syrup as already described (Aufauvre et al. 2014). Cohorts of *ca*. 50 honeybees were chronically exposed *per os* for 21 days to none or to increasing concentrations of DMCH or OO in the sucrose syrup (0, 0.25, 0.5, 0.75, 1.0, 1.25, 1.7 and 3.4 *μ*g/L). All treatments were performed in four replicates. Dead bees were removed and counted daily and the honeybee proportion of survival was assessed (Aufauvre et al. 2014). Survival data are available on request. Honeybee survival was significantly reduced in groups exposed to OO at 1.25 and 3.4 *μ*g/L and for the groups exposed to DMCH at 0.75 and 3.4 *μ*g/L (generalized Wilcoxon test, *p*<*0*.*05*, data not shown). For OO at 1.25 *μ*g/L and DMCH at 0.75 *μ*g/L, the differences were mainly drived by one of the four replicates, suggesting an artifactual or cohort-dependent effect. At a concentration of 3.4 *μ*g/L, both VOCs became clearly toxic for honeybees. However, both VOCs did not significantly affect the proportion of survival at the end of the experiment, whatever their concentration (Dunn test, *p*>*0*.*05*). The two VOCs had transitory toxicity, that was no more visible after 21 days. This could be due to a variability of sensitivity between honeybees.

The mass of consumed sucrose was also monitored daily. OO did not affect sucrose consumption while a significant decrease of 10.2 ± 4%, was observed for DMCH (Dunn test, *p*>*0*.*05*), suggesting a potential repellent effect. Some VOCs are known to exert such repulsion for contaminated food in insects (Leroy et al. 2011; Saha and Chandran 2017). The data led us to select OO for a further analysis since it appeared poorly toxic and not repellent at the tested conditions.

A second experiment was then performed to evaluate the effect of OO on the survival of bees chronically exposed to low doses of fipronil. Cohorts of *ca*. 70 encaged workers were chronically exposed *per os* for 21 days to 0.5 or 1.0 *μ*g of fipronil per L of sucrose syrup (Aufauvre et al. 2014) and/or to four concentrations of OO (0.5, 1.0, 1.7 and 3.4 *μ*g/L). All treatments were performed in four replicates. The honeybee proportion of survival was recorded (Fig. 1). The lower concentration of fipronil reduced the survival of honeybees (generalized Wilcoxon test, *p*<0,05) but with variations among replicates, with two out of four that showed a significant reduction of survival. The fipronil high concentration significantly reduced the survival for all replicates. After 21 days, the proportions of survival were of 26 ± 8% for control, 16 ± 16% for fipronil at 0.5 *μ*g/L, and 3,2 ± 4,3% for fipronil at 1 *μ*g/L.

**Figure 1.**
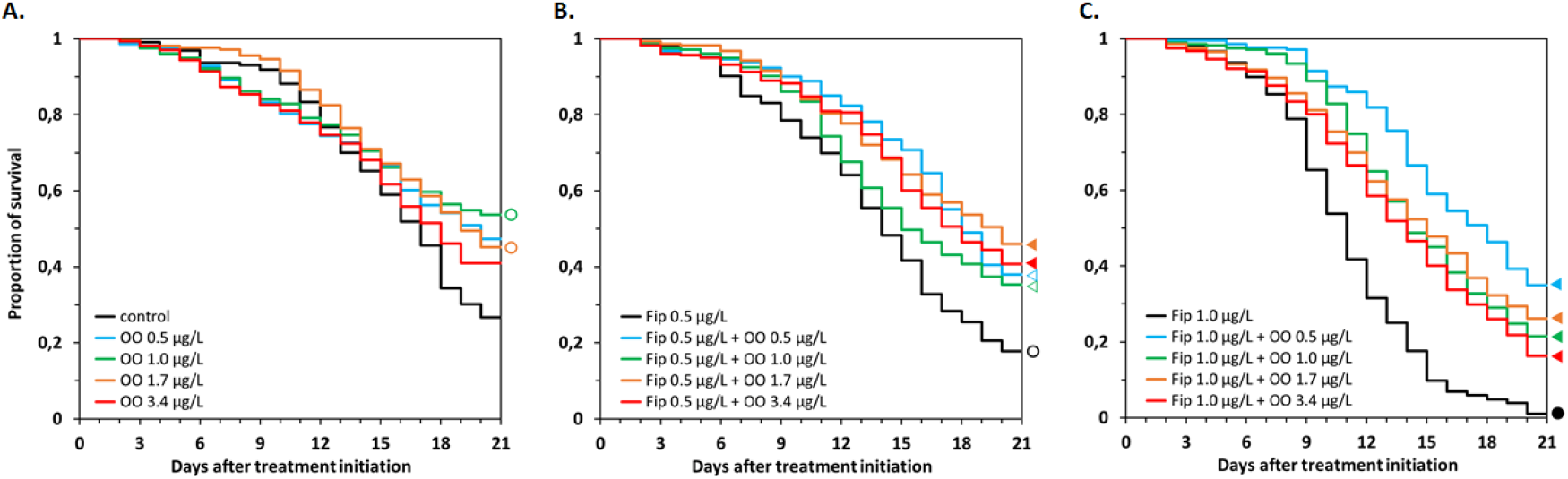
Survival of honeybees chronically exposed to fipronil (Fip) and to 1-octen-3-ol (OO). Workers were orally exposed to none and four concentrations of OO, given in *μ*g per L of sucrose syrup **(A)**, to fipronil at 0.5 *μ*g/L and OO **(B)** and to fipronil at 1.0 *μ*g/L and OO **(C)**. The proportions of survival represent the pools of four replicates of cohorts of *ca*. 70 bees. A dot represents a significant difference (Wilcoxon test, *p*<*0*.*05*) compared to the control condition, due to 2 out of 4 (○) or to all replicates (•). A triangle represents a significant difference compared to the fipronil-alone condition, due to 2 out of 4 (Δ) or to all replicates (◂).

For honeybees co-exposed to the low fipronil concentration, a significant increase in survival was observed with the four tested OO concentrations compared to those exposed to fipronil only (Fig. 1B). For the two lower concentrations of OO, only two out of four replicates responded significantly while all replicates were significant for the two higher concentrations. A higher increase in survival was observed for honeybees co-exposed to OO and to the highest fipronil concentration (Fig. 1C), with all replicates significant. These differences were also observed when comparing the honeybee survival after 21 days (Dune test, *p*<*0*.*05*), with an average increase in survival of 25.7 ± 2.9% and 21.3 ± 9.4% for groups co-exposed to OO and fipronil at 0.5 and 1 *μ*g/L respectively. There was no significant difference between the survival of honeybees co-exposed to OO and fipronil at 0.5 *μ*g/L and those exposed to the corresponding OO concentration alone. In contrast, the mortality was higher in the presence of 1 *μ*g/L fipronil suggesting that the survival was only partly restored by OO. In conclusion, OO seemed to have a beneficial effect on the survival of *A. mellifera* chronically exposed to fipronil.

No difference in the daily sucrose consumption was observed between the control group and the fipronil-exposed groups (Dunn test, *p*>*0*.*05*). However, the co-exposed groups consumed significantly less sucrose than groups exposed to fipronil alone (Dunn test, *p*<*0*.*05*), representing a decrease of 10.5 ± 3.9% and 16.7 ± 6.2% for the groups co-exposed with fipronil at 0.5 *μ*g/L and 1 *μ*g/L respectively. This change in feeding behavior may have reduced both insecticide and OO intake, partly explaining a higher survival. Yet, the cumulative consumption did not differ at the end of the experiment (Mann-Whitney, *p*>*0*.*05*). Thus, the improvement of survival was probably not only due to a change of the feeding behavior.

Hypotheses may be proposed to explain the observed beneficial effect of OO on the survival of *A. mellifera* chronically exposed to fipronil. OO may exert a direct action on GABA-R, as demonstrated in mammals (Johnston, 2006; Köksal, 2015). It would be interesting to determine whether OO actually interacts with GABA-Rs in *A. mellifera* and whether its action counteracts that of fipronil. OO could also have hormetic effect, *i*.*e*. it may induce an overall improvement in the honeybee capacity to respond to insecticide stress. Indeed, an improved survival was observed in some cohorts of honeybees exposed to OO only (Fig. 1A). Thus, a better survival could have been an indirect effect of the VOC. Fipronil exerts a variety of sublethal effects (Pisa et al. 2015), that could have been buffered by OO. In order to detect potential involved mechanisms, the expression of genes involved in oxidative stress management, or in xenobiotic and GABA metabolism (catalase, superoxide dismutase, glutathione transferase S1, GABA aminotransferase, glutamate decarboxylase) was analysed by RT-qPCR in honeybees head after two, four and seven days of exposure. No difference in expression was observed whatever the considered experimental condition and time point. At last, the honeybee survival was much lower in the second experiment (21 ± 13 % for the control condition), where OO showed positive effect but no toxicity, than in the first experiment (86 ± 7 % for the control condition), where OO showed toxicity but no positive effect (data not shown). This suggests that the positive or negative action of OO varies with the general health status of the honeybee. OO may have a beneficial effect only for honeybees with a weak probability of survival while showing potential toxicity for healthy honeybees.

In conclusion, while our results give hint on a possible beneficial effect of OO, more data ought to be collected before concluding towards a protective effect of OO against fipronil. It is therefore essential to repeat the experiments with more replicates from distinct colonies with various environmental (*e*.*g*. landscapes with different resources and contaminants) and genetic backgrounds and by treating larger cohorts and/or whole colonies.

## Statements & Declarations Funding

This research was funded by the UMR CNRS 6023 LMGE and the AURA Environmental Research Federation.

## Conflicts of interest/Competing interests

The authors declare they have no competing interest and no relevant financial or non-financial interests to disclose.

## Ethics approval/Declarations

No ethical approval was required for the study.

## Consent to participate

Not applicable

## Consent for publication

Not applicable

## Availability of data and material

Survival data are available on request

## Code availability

Not applicable

## Author’s contribution

VF, TS, FD and PB contributed to the study conception and design. VF, TS and MS performed the experiments. VF, TS, MS and NB analyzed the data. The first draft of the manuscript was written by VF and TS. All authors commented and reviewed the manuscript. All authors read and approved the final manuscript.

## Notes

### Competing Interest Statement

The authors have declared no competing interest.

